# *Mutator* foci are regulated by developmental stage, RNA, and the germline cell cycle in *Caenorhabditis elegans*

**DOI:** 10.1101/2020.06.17.158055

**Authors:** Celja J. Uebel, Dana Agbede, Dylan C. Wallis, Carolyn M. Phillips

## Abstract

RNA interference is a crucial gene regulatory mechanism in *Caenorhabditis elegans*. Phase-separated perinuclear germline compartments called *Mutator* foci are a key element of RNAi, ensuring robust gene silencing and transgenerational epigenetic inheritance. Despite their importance, *Mutator* foci regulation is not well understood, and observations of *Mutator* foci have been largely limited to adult hermaphrodite germlines. Here we reveal that punctate *Mutator* foci arise in the progenitor germ cells of early embryos and persist throughout all larval stages. They are additionally present throughout the male germline and in the cytoplasm of post-meiotic spermatids, suggestive of a role in paternal epigenetic inheritance. In the adult germline, transcriptional inhibition results in a pachytene-specific loss of *Mutator* foci, indicating that *Mutator* foci are partially reliant on RNA for their stability. Finally, we demonstrate that *Mutator* foci intensity is modulated by the stage of the germline cell cycle and specifically, that *Mutator* foci are brightest and most robust in the mitotic cells, transition zone, and late pachytene of adult germlines. Thus, our data defines several new factors that modulate *Mutator* foci morphology which may ultimately have implications for efficacy of RNAi in certain cell stages or environments.

## INTRODUCTION

RNA interference (RNAi) is an evolutionarily conserved strategy to ensure proper gene expression across a wide range of eukaryotes (Fire *et al*. 1998; Shabalina and Koonin 2008). The effectors of RNAi are members of the Argonaute protein family, which bind small regulatory RNAs ranging from 18-30 nucleotides in length. Together these components form the RNA Induced Silencing Complex (RISC), which targets fully or partially complementary transcripts of exogenous or endogenous origin through direct cleavage, recruitment of exonucleases, or repression of translational complexes (Hutvagner and Simard 2008; Wu and Belasco 2008). By these means, small RNAs play key roles in development, fertility, chromosome segregation, and defense against foreign genetic elements such as transposons and viruses.

In *C. elegans*, RNA silencing is mediated by a network of ~27 Argonaute proteins associated with three distinct classes of small RNAs: micro-RNAs (miRNAs), Piwi interacting-RNAs (piRNAs), and small interfering RNAs (siRNAs). *C. elegans* siRNAs are categorized in two distinct groups: primary siRNAs and secondary siRNAs. Primary siRNAs are cleaved by Dicer from double-stranded RNA and are bound by primary Argonaute proteins like RDE-1 and ERGO-1, whereas secondary siRNAs are produced from primary siRNA-targeted or piRNA-targeted templates by the RNA-dependent RNA polymerases (RdRPs), RRF-1 and EGO-1, and are bound by the worm-specific Argonaute (WAGO) clade of Argonaute proteins (Yigit *et al*. 2006; Aoki *et al*. 2007; Pak and Fire 2007; Sijen *et al*. 2007; Gu *et al*. 2009; Vasale *et al*. 2010). Secondary siRNAs are crucial for signal amplification and transgenerational silencing; mutant animals that fail to produce secondary siRNAs display multiple defects including failure to respond to RNAi, temperature-sensitive sterility, and transposon mobilization (Ketting *et al*. 1999; Tabara *et al*. 1999; Vastenhouw *et al*. 2003; Zhang *et al*. 2011).

Secondary siRNA amplification primarily occurs in the *mutator* complex, which forms perinuclear foci, known as *Mutator* foci, in *C. elegans* germ cells (Phillips *et al*. 2012). These *Mutator* foci are nucleated by MUT-16, which directly interacts with the RdRP RRF-1, and other key siRNA biogenesis proteins to assemble *mutator* complexes at these sites (Phillips *et al*. 2012; Uebel *et al*. 2018). The C-terminal region of MUT-16, which contains regions of intrinsic disorder, is both necessary and sufficient for *Mutator* foci formation. Furthermore, *Mutator* foci behave as phase-separated condensates, with liquid-like properties such as rapid recovery after photobleaching, temperature and concentration dependent condensation, and disruption after perturbation of weak hydrophobic interactions (Uebel *et al*. 2018). Additionally, *Mutator* foci are adjacent to P granules, which are phase-separated mRNA surveillance centers important for maintenance of the germ cell fate and fertility (Pitt *et al*. 2000; Brangwynne *et al*. 2009; Sheth *et al*. 2010; Updike *et al*. 2014; Campbell and Updike 2015; Knutson *et al*. 2017). Because P granules also contain proteins associated with the small RNA pathways, including both primary and secondary Argonaute proteins (Wang and Reinke 2008; Claycomb *et al*. 2009; Gu *et al*. 2009; Conine *et al*. 2010), we hypothesize that P granules and *Mutator* foci interact at the nuclear periphery to coordinate small RNA silencing of nascent germline transcripts.

Though P granule regulation and morphology has been extensively studied in all stages of *C. elegans* development (Strome and Wood 1982; Updike and Strome 2009), investigation of *Mutator* foci is primarily limited to observations within the adult hermaphrodite germline (Phillips *et al*. 2012; Uebel *et al*. 2018). Here, we examine *Mutator* foci throughout embryonic, larval, male, and hermaphrodite germline development and begin to assess factors that regulate or influence *Mutator* foci morphology. We determine that *Mutator* foci first appear as bright, distinct puncta in the Z2/Z3 progenitor germ cells (PGCs), and that these foci persist throughout all subsequent developmental stages. We then demonstrate that *Mutator* foci are present in the male germline throughout spermatogenesis, and that MUT-16 is sequestered into the cytoplasm of post-meiotic spermatids. While probing potential regulatory mechanisms for *Mutator* foci, we discover that these phase-separated compartments are at least partially dependent on RNA for their stability in pachytene region of the gonad. Finally, we find that *Mutator* foci are largest and most robust in the mitotic cells, the transition zone, and the late pachytene of adult germlines. By RNAi of key germline development proteins, we discover that the mitotic cell stage, but not the transition zone, is a determinant of robust *Mutator* foci. Thus, through these observations, we better define *Mutator* foci in all developmental stages and probe the regulatory factors that influence morphology of this secondary siRNA amplification center.

## MATERIALS AND METHODS

### *C. elegans* Strains

Worms were grown at 20°C according to standard conditions (S. Brenner 1974). Strains used in this study include:

**USC1266** *mut-16(cmp41[mut-16::mCherry::2xHA + loxP]) I; pgl-1(sam33[pgl-1::gfp::3xFLAG]) IV*
**USC717** *mut-16(cmp3[mut-16::gfp::3xFLAG + loxP]) I* (Uebel *et al*. 2018)
**USC1385** *mut-16(cmp3[mut-16::gfp::3xFLAG + loxP]) I; him-8(tm611) IV*

USC1266 was created by crossing DUP75 (*pgl-1(sam33[pgl-1::gfp::3xFLAG]*) (Andralojc *et al*. 2017) and USC896 (*mut-16(cmp41 [mut-16::mCherry::2xHA + loxP]*) (Uebel *et al*. 2018; Nguyen and Phillips 2020). USC1385 was created by crossing USC717 (*mut-16(cmp3[mut-16::gfp::3xFLAG + loxP])*) (Uebel *et al*. 2018) and CA257 (*him-8(tm611)*) (Phillips *et al*. 2005).

### Antibody staining and microscopy

For fixed embryo imaging, gravid adult *mut-16::mCherry::2xHA; pgl-1::gfp::3xFLAG* animals were placed in 4μL water on SuperFrost slides and burst to release embryos by application of a glass coverslip. Slides were placed on a dry-ice cooled aluminum block for freeze-crack permeabilization. After coverslip removal, slides were fixed in 100% ice-cold Methanol for 15 minutes and washed three times in 1xPBST for 5 minutes each. DAPI was added to the first wash. Embryos were mounted in 10μL NPG-Glycerol mounting medium and imaged (Phillips *et al*. 2009). Embryos were staged by approximate cell counts and position of the Z2/Z3 PGCs. To avoid fluorescent artifacts, no antibody staining was used, as the proteins of interest were fluorescently tagged via CRISPR at their endogenous loci. To avoid FRET activation and channel bleed-through, twenty 0.2 μm Z-stacks were captured first with 542 nm (red), followed by 475 nm (green) and 390 nm (blue) excitation.

For live imaging, undissected larva, hermaphrodites, or adult males were mounted in M9 containing <1% sodium azide to inhibit movement or in M9 with no sodium azide, and images were collected first with 542 nm excitation followed by 475 nm laser excitation. Larval images were compiled from 10 Z-stacks and staged by gonad morphology. Male germline live images were compiled from 20 Z-stacks. Adult hermaphrodite germline images were compiled from five Z-stacks.

For immunostained germlines, adult males or hermaphrodites were dissected in egg buffer containing 0.1%Tween-20 and fixed in 1% formaldehyde as described (Phillips *et al*. 2009). *mut-16::GFP::3xFLAG; him-8(tm611)* germlines were immunostained with 1:500 rabbit anti-GFP (Thermo Fisher A-11122), and all other germlines were stained with 1:2000 mouse anti-FLAG (Sigma F1804) and 1:250 rabbit anti-SYP-1 (Macqueen *et al*. 2002). Fluorescent Alexa-Fluor secondary antibodies were used at 1:1000 (Thermo Fisher).

All imaging was performed on a DeltaVision Elite (GE Healthcare) microscope using a 60x N.A. 1.42 oil-immersion objective. For all images, the 0.2 μm Z-stacks were compiled as maximum intensity projections and pseudo-colored using Adobe Photoshop. Image brightness and contrast were adjusted for clarity.

### Transcriptional Inhibition

Anterior gonads of young adult (~24 hours post L4) hermaphrodites were microinjected with 200 μg/mL α-amanitin until the solution flowed around the germline bend, as previously described (Uebel and Phillips 2019). A vehicle control injection of RNase free water did not disrupt foci, nor were foci disrupted in the uninjected posterior gonad. The observed results were reproducible despite slight variation in microinjected volume and gonad integrity. Due to injection and imaging time requirements, animals were imaged at three, five, and seven hours ±15 minutes post-injection. At least three animals were imaged for each time point.

### RNA interference

For all RNAi experiments, sequence-confirmed RNAi clones *gld-1, gld-2, chk-2, atx-2, ama-1* and L4440 (control) in HT115 (DE3) bacteria were grown overnight at 37°C to maximum density. Cultures were concentrated 10-fold and 100μL was plated on NGM plates with 5 mM IPTG and 100 μg/mL Ampicillin. For *gld-1, gld-2* double RNAi, the concentrated cell cultures were combined at equal volume before plating. Plates were stored at room temperature for at least 24 hours to allow for IPTG induction. Synchronized L1 worms were then plated and grown at 20°C for 70 hours before dissection and imaging, unless otherwise stated. A minimum of five animals for each clone in two independent experiments were analyzed to confirm RNAi efficiency and phenotype.

### Quantification of *Mutator* foci intensity

Gonads were divided by length into 10 equal regions ending at the first single-file diplotene/diakinesis nucleus. Raw TIFs were loaded into FIJI and backgrounds were subtracted with a 50-pixel rolling ball radius. Each image was then thresholded identically, creating a binary image displaying only above-threshold foci. Each region was processed using the “analyze particles” function and the resulting particle quantification data was collected and graphed in Excel.

### Data Availability

All strains are available either at the Caenorhabditis Genetics Center (CGC) or upon request from the Phillips lab. The authors affirm that all data necessary for confirming the conclusions of the article are present within the article, figures, and tables.

## RESULTS

### Robust *Mutator* foci first appear in the Z2/Z3 progenitor germ cells in *C. elegans* embryos

Current characterization of *Mutator* foci is largely restricted to observations within the adult germline, without consideration for earlier stages of development. *C. elegans* embryos undergo invariant cell divisions, the first of which gives rise to the asymmetric AB and P_1_ blastomeres. The P_1_ cell subsequently produces the P_2_, P_3_, and P_4_ cells, the latter of which divides around the 100-cell stage to form the Z2 and Z3 PGCs, which undergo no further divisions until hatching and feeding. After hatching, the PCGs eventually give rise to the ~2,000 cells that comprise the adult germline. Recent imaging by Ouyang *et al*. (2019) observes punctate *Mutator* foci in the PGCs of later comma-stage embryos (>550 cells), which describes the onset of elongation around 400 minutes post fertilization. To determine the earliest formation of *Mutator* foci, defined here as distinctly punctate MUT-16 fluorescence, we imaged endogenous MUT-16::mCherry and PGL-1::GFP in fixed embryos at representative developmental stages. At the 2-cell stage, P granules segregate to the cytoplasm of the P_1_ blastomere, while MUT-16 predominantly appears as diffuse cytoplasmic signal in both AB and P_1_ cells (Figure 1A and A’). At the 16-cell stage, P granules begin to associate with the nuclear periphery of the P4 cell (Updike and Strome 2010), yet MUT-16 remains diffusely cytoplasmic in all cells of the embryo (Figure 1B and B’). As the embryos reach the 30-cell stage, coinciding with the beginning of gastrulation (Sulston *et al*. 1983), MUT-16 continues to be expressed throughout the cytoplasm of all embryonic cells (Figure 1C), though we sometimes observe faint punctate *Mutator* foci adjacent to the large, perinuclear P granules (Figure 1C’ and Supplemental Figure 1). At this stage, any PGL-1 remaining in the somatic blastomeres is eliminated via autophagy, temporarily creating small PGL-1 foci throughout the embryo (Hird *et al*. 1996; Zhang *et al*. 2009), yet diffuse MUT-16 expression in the cytoplasm is unchanged. Interestingly, in addition to the cytoplasmic MUT-16 signal in the 100-cell stage, we consistently observe punctate *Mutator* foci surrounding the newly formed Z2 and Z3 PGCs (Figure 1D). The distinct foci are adjacent to P granules and at the nuclear periphery, comparable to *Mutator* foci localization in adult germ cells. Since we observe foci in the 300-cell stage, and Ouyang *et al*. (2019) observes foci in the later comma stage, *Mutator* foci appear to persist throughout later stages of embryonic development (Figure 1E and E’). Thus, our imaging reveals that diffuse cytoplasmic MUT-16 is present in all embryonic cells at all stages of embryonic development, and punctate *Mutator* foci can form as early as the 30-cell stage, but are most consistent and robust in the Z2/Z3 PGCs.

**Figure 1.**
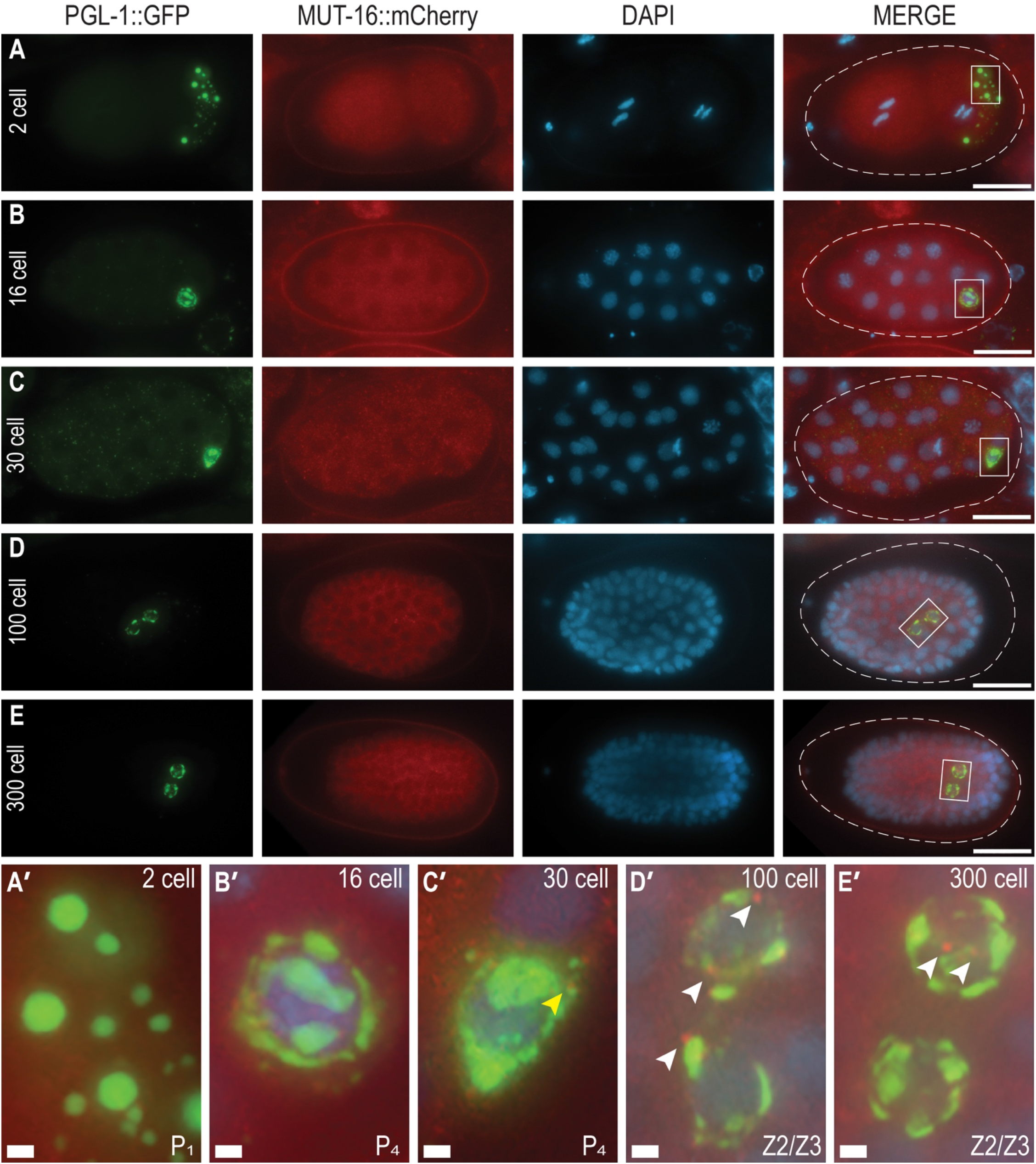
Distinct *Mutator* foci appear in the Z2/Z3 progenitor germ cells. (A-E) Distribution of PGL-1::GFP (green) and MUT-16::mCherry (red) in methanol-fixed embryos of representative stages. DNA is stained with DAPI (blue) for visualization. Scale bars, 15μm. (A’-E’) Enlarged insets of boxed areas from merged embryos in A-E. Small MUT-16 puncta are visible in the 30-cell stage (yellow arrowhead). Distinct MUT-16 foci are consistently visible in Z2/Z3 cells of both 100- and 300-cell embryos (white arrowheads). Scale bars, 1μm.

### *Mutator* foci persist in all larval stages of *C. elegans*

To expand on our characterization of *Mutator* foci throughout development, we examined MUT-16 and PGL-1 expression during all larval stages, including the “survival-state” dauer stage induced by starvation (Cassada and Russell 1975). In fed L1 larva, the PGCs are easily identified by perinuclear P granules and, despite intestinal autofluorescence, MUT-16::mCherry is clearly visible in punctate *Mutator* foci (Figure 2A). *Mutator* foci are present in both early and late L2 stages (Figure 2B-C). Additionally, *Mutator* foci are visible in the L2d dauer larva, whose germlines are developmentally quiescent (Figure 2D). Finally, *Mutator* foci are present in the early L3 developmental stage, at which point two “arms” of the gonad begin to form and branch from the central somatic gonad primordium (Figure 2E). Similar to MUT-16 expression in adults, larval germ cells also show faint cytoplasmic expression and MUT-16 exclusion from the nucleus. Since *Mutator* foci have been observed previously in L4 and adult stages (Phillips *et al*. 2012), we can conclude that *Mutator* foci are present across all stages of larval development.

**Figure 2.**
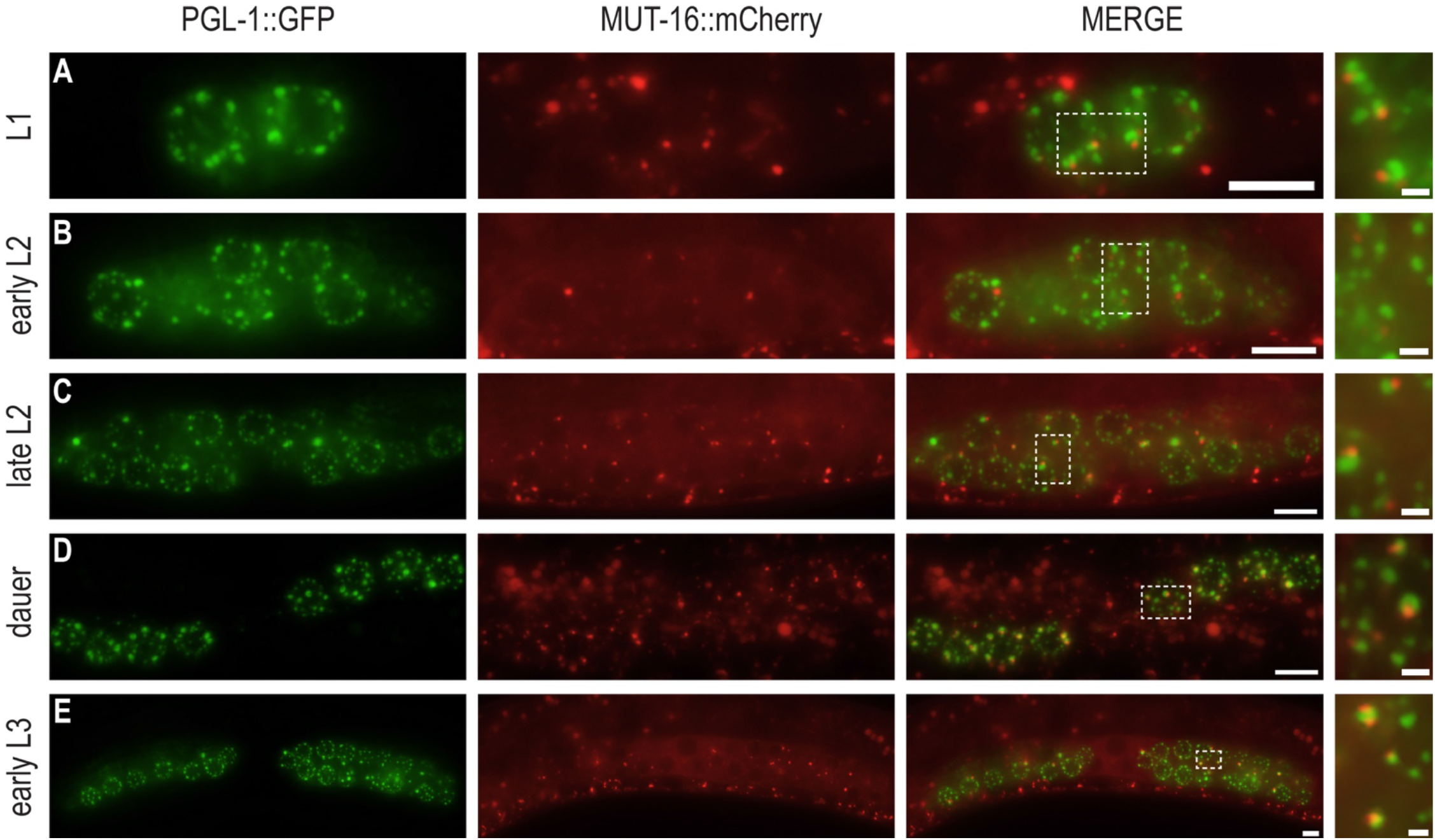
*Mutator* foci are present in all larval stages. (A-E) Live imaging of *mut-16::mCherry; pgl-1::gfp* expression in representative larval stages. PGL-1::GFP granules (green) associate with germ cells in the developing gonad in all larval stages. MUT-16::mCherry foci (red) are also present. Merged panels and enlarged insets (dashed region) show PGL-1 granules and MUT-16 foci interacting at the nuclear periphery. Merge scale bars, 5μm. Inset scale bars, 1μm.

### *Mutator* foci localizes throughout spermatogenesis and MUT-16 is deposited in the cytoplasm of spermatids

In addition to an emphasis on adult germlines, current studies of *Mutator* foci rely almost exclusively on hermaphrodite germlines for characterization. To fully characterize *Mutator* foci in male germlines, we live imaged *mut-16::mCherry; pgl-1::gfp* males. *Mutator* foci appear similar to hermaphrodite germlines throughout the mitotic tip, the transition zone, and the pachytene region (Figure 3A). However, we observe a divergence in the localization patterns of P granules and *Mutator* foci in spermatogenesis. Previous literature reports that PGL-1 and PGL-3 disassemble in late spermatogenesis, though the P granule component GLH-1 persists until eventual segregation into residual bodies of budding spermatids (Gruidl *et al*. 1996; Amiri *et al*. 2001; Updike and Strome 2010). While we observe PGL-1 disassembly in late spermatogenesis, we continue to see punctate *Mutator* foci around nuclei with no PGL-1 signal (Figure 3A). *Mutator* foci were previously observed to localize to the nuclear periphery despite lack of detectable PGL-1 via a *glh-1/glh-4* RNAi knockdown (Phillips *et al*. 2012), and our findings support the independent localization of *Mutator* foci.

**Figure 3.**
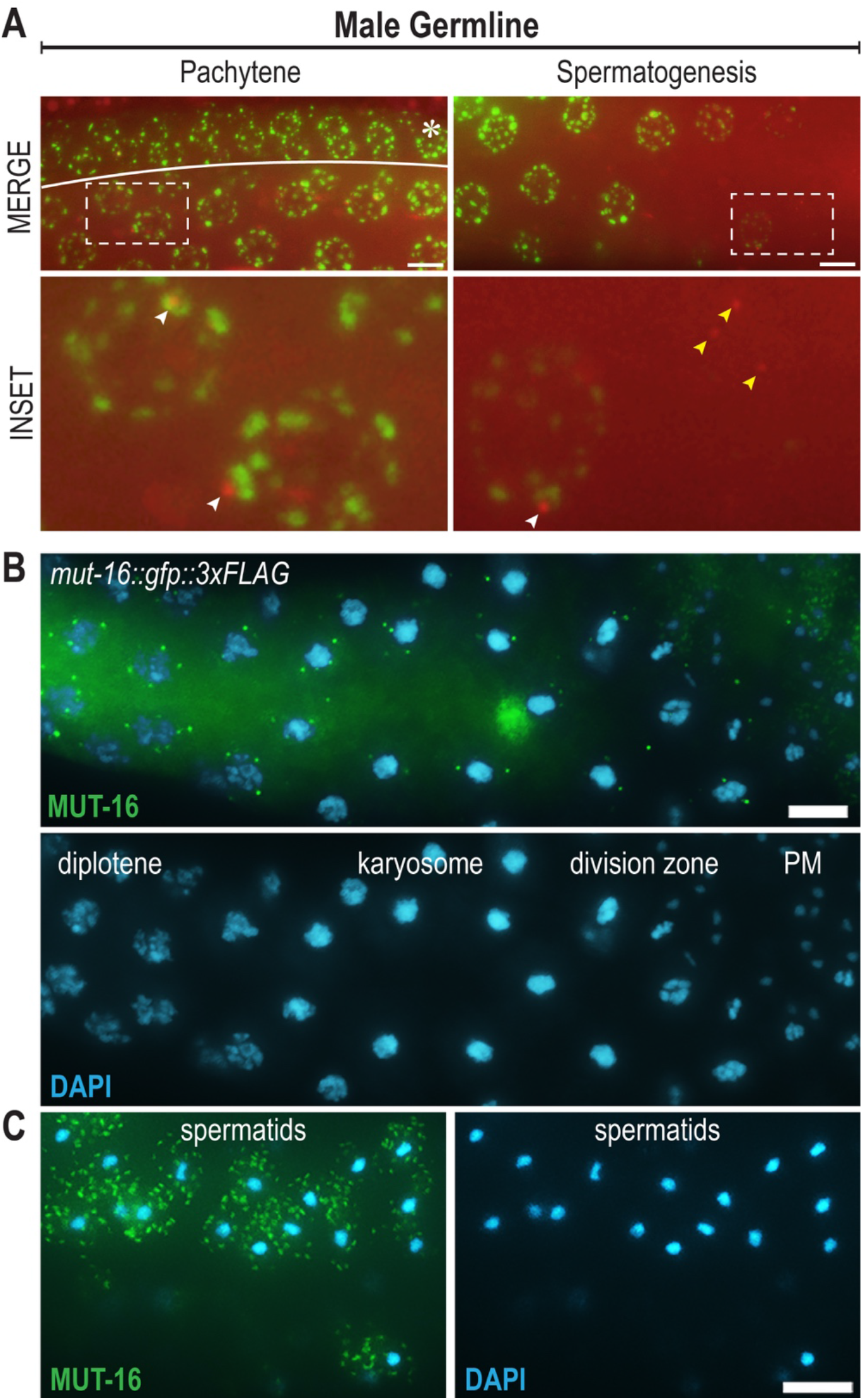
*Mutator* foci are present throughout spermatogenesis and are found in the cytoplasm of spermatids. (A) Live images of undissected *mut-16::mCherry; pgl-1::gfp* male germlines. The distal region (asterisks) is folded over the pachytene region. Inset (dashed box) is enlarged below to show MUT-16 foci (red) and PGL-1 granule (green) interactions (white arrows). In late spermatogenesis, PGL-1::GFP signal becomes undetectable, while punctate MUT-16 foci remain visible (yellow arrows). (B) MUT-16 foci (anti-FLAG, green) are present through meiotic division in dissected and immunostained male germlines. Chromosome morphology is visualized by DAPI (blue) and distinguishes diplotene nuclei, karyosome formation, the division zone, or post-meiotic (PM) nuclei. (C) MUT-16 localizes in a granular pattern within the cytoplasm of post-meiotic spermatids. All scale bars, 5μm.

To visualize *Mutator* foci and corresponding nuclei more clearly throughout spermatogenesis, we dissected and fixed *mut-16::gfp* male germlines. Using DAPI-stained DNA as a guide for morphological identification of chromatin state, we observe punctate *Mutator* foci throughout the condensation zone and into the division zone (Figure 3B). *Mutator* foci are present around nuclei in the karyosome stage, a poorly understood state of nuclear compaction thought to promote chromosome organization prior to meiotic division (Shakes *et al*. 2009). Because chromosomes are highly condensed in the karyosome stage and the nuclear envelope size remains unchanged, *Mutator* foci appear farther away from the DAPI-stained bodies. Additionally, during late diakinesis and metaphase, the nuclear envelope begins to break down, which may explain why some punctate *Mutator* foci in the division zone are no longer associated with DAPI-stained nuclei (Figure 3B).

Unexpectedly, we discovered MUT-16::GFP signal in the cytoplasm of post-meiotic spermatids in a unique granular pattern reminiscent of WAGO-1 localization in spermatids (Conine *et al*. 2010) (Figure 3C). The presence of MUT-16 in spermatids carries implications for paternal inheritance of *Mutator* components and paternal deposition of WAGO class 22G-RNAs, which have been shown to rescue fertility of piRNA-depleted RNAi defective hermaphrodites (Phillips *et al*. 2015).

### RNA influences *Mutator* foci stability and morphology

Many phase-separated condensates are comprised of multivalent interactions between RNA and proteins, often referred to as ribonucleoprotein (RNP) granules. Concentration, secondary structure, and RNA length have been shown to regulate size, interaction, and formation of RNP granules both *in vivo* and *in vitro* (Langdon *et al*. 2018; Garcia-Jove Navarro *et al*. 2019). Previously work demonstrated that the presence of RNA is integral to sustaining P granule integrity *in vivo* (Sheth *et al*. 2010). Five hours after gonadal microinjection of a potent RNA Polymerase II transcriptional inhibitor, α-amanitin, P granules disappeared in a pachytene-specific manner. To determine if *Mutator* foci also require RNA for their stability, we microinjected 200 μg/mL α-amanitin into the gonads of adults expressing fluorescently-tagged MUT-16 and PGL-1. We found that both P granules and *Mutator* foci began to disperse by three hours, forming fewer foci in both the mid and late-pachytene regions (Figure 4A and 4B). In late pachytene, P granules appeared rounder and more detached from the nuclear pore environment (Figure 4B). *Mutator* foci remain adjacent to P granules despite the overall reduction in foci number.

**Figure 4.**
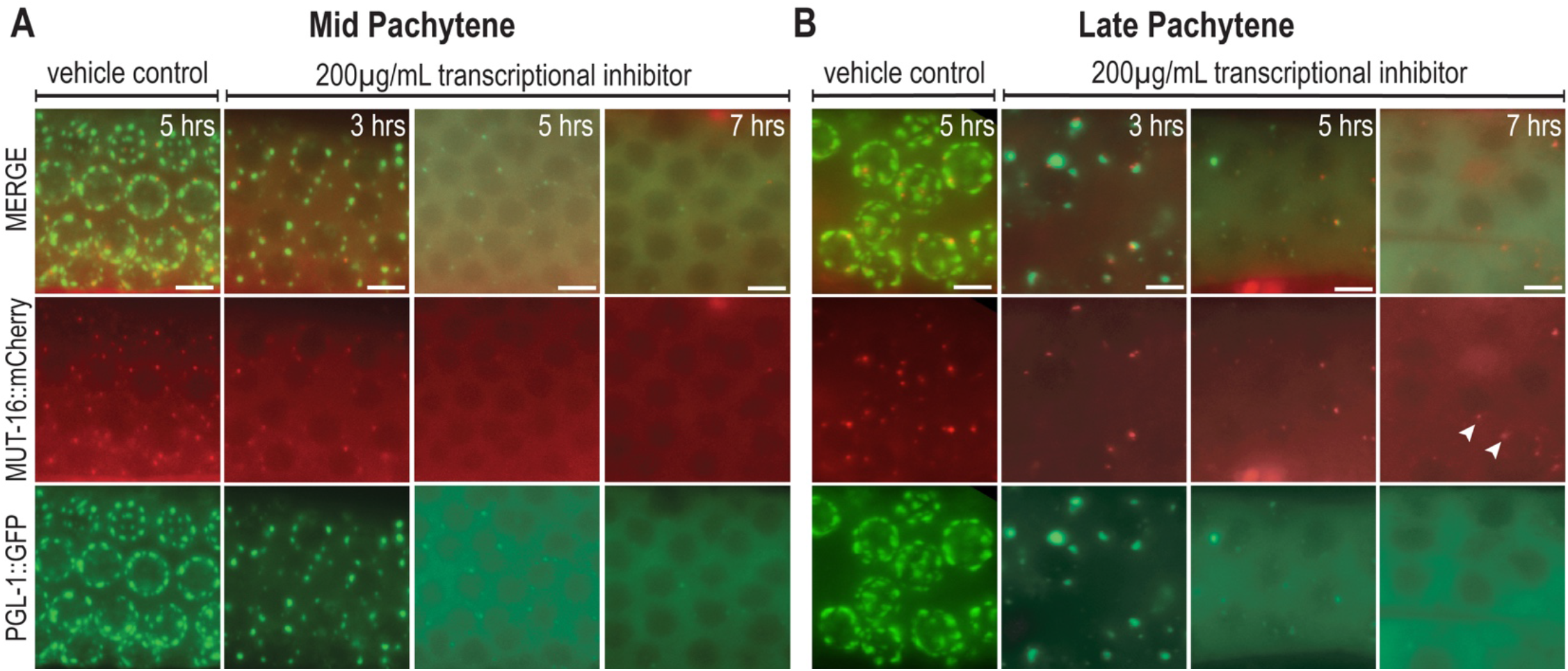
*Mutator* foci integrity partially relies on RNA. Germlines of young adult *mut-16::mCherry; pgl-1::gfp* animals were injected with either a vehicle control or 200 μg/mL α-amanitin, a transcriptional inhibitor. (A) Live images of the mid-pachytene region at three, five, or seven hours post injection. MUT-16 foci (red) and PGL-1 granules (green) are unperturbed in vehicle control injections but dissipate with increasing severity over time. (B) Live images of the late-pachytene region also reveal an increasing severity of dissipation over time. At seven hours, some MUT-16 foci remain (white arrows) though PGL-1 granules are absent. Scale bars, 5μm.

By five and seven hours post α-amanitin injection, P granules and *Mutator* foci are nearly completely dissolved in the mid-pachytene region (Figure 4A). In late pachytene, P granules and *Mutator* foci also dissipate with increasing severity (Figure 4B). Interestingly however, we consistently observe faint *Mutator* foci remaining at seven hours, despite no detectible PGL-1 foci. This suggests that *Mutator* foci may only partially rely on RNA for stability.

To further explore the effect of mRNA loss on *Mutator* foci, we placed L4 animals on RNAi targeting *ama-1*, the largest subunit of RNA polymerase II, required for mRNA transcription (Bird and Riddle 1989). Phenotypes from some RNAi can be first observed after 24 hours of feeding (Kamath *et al*. 2001). We did not observe any noticeable perturbation of either P granules or *Mutator* foci at 24 hours on *ama-1* RNAi, but after 30 hours, PGL-1 granules in the mid-pachytene were faint and fluorescent signal was largely cytoplasmic (Supplemental Figure 2). Similarly, MUT-16 foci dissipated after 30 hours on *ama-1* RNAi. Though it is possible that the inhibition of transcription resulted in reduction of PGL-1, MUT-16, or other major protein constituents of these granules, the dispersed cytoplasmic signal of PGL-1 and MUT-16 is more suggestive of foci loss due to lack of mRNA than due to lack of protein components. This data supports our hypothesis that *Mutator* foci at least partially rely on RNA for foci integrity.

### *Mutator* foci intensity is dictated by stage of germ cell progression

To examine other elements of *Mutator* foci regulation, we directed our attention to germ cell stage and morphology. The *C. elegans* germline is a syncytium of cells organized in two branching “arms” extending from a central somatic uterus. The distal tip of each arm contains actively dividing mitotic cells that progress through meiotic S phase before transitioning to the leptotene/zygotene stage of meiosis. In this transition zone, nuclei appear crescent-shaped as chromosomes polarize to find homologs, a trait that is readily apparent by DAPI staining. The transition zone is followed by the early, mid, and late pachytene stage, where crossing over of homologous chromosomes occurs. Cells then progress proximally through the diplotene and diakinesis stages, where chromosomes condense and the nuclear envelope begins to break down. By live and immunofluorescent imaging of endogenously tagged MUT-16, we noticed the early- and mid-pachytene regions of adult germlines contain dim *Mutator* foci, whereas the mitotic tip, transition zone, and late pachytene regions have large, bright *Mutator* foci (Figure 5A). We refer to this quality of brightness as “*Mutator* foci intensity”. To quantify *Mutator* foci intensity, we divided each gonad into 10 equal sections and performed a granule count on images processed with a uniform threshold (Supplemental Figure 3). In this manner, dim foci fell below the threshold and only bright foci were counted, allowing us to determine the distribution of the brightest foci throughout the germline. To help delineate meiotic stage, we utilized a central component of the synaptonemal complex, SYP-1, which loads during the transition zone and is present on synapsed chromosomes through the pachytene region (MacQueen 2002). We discovered that the first two regions of the gonad, which corresponds to the mitotic zone, contains high percentages of above-threshold *Mutator* foci (Figure 5B, regions 1-2). However, *Mutator* foci intensity peaks in a single region coinciding with the onset of the transition zone (Figure 5B, region 3). This peak in *Mutator* foci intensity is followed by a dramatic reduction in intensity in the regions corresponding to early and mid pachytene (Figure 5B, regions 4-7). Consistent with our qualitative observations, the foci-depleted region is followed by an increase in foci intensity in regions corresponding to the late pachytene (Figure 5B, regions 8-10). Our quantification shows *Mutator* foci intensity fluctuates in a consistent bright-dim-bright pattern within adult wild-type germlines.

**Figure 5.**
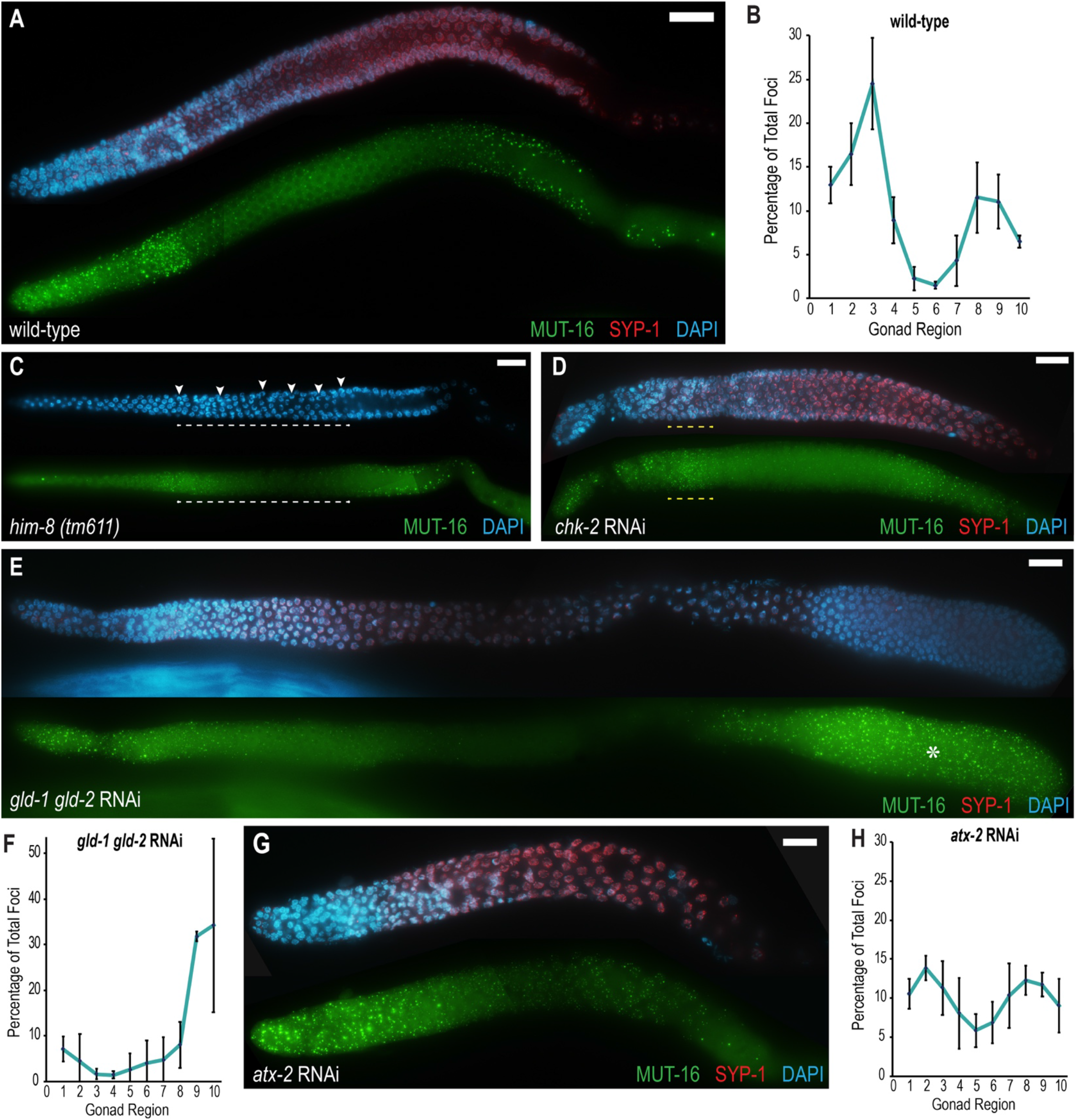
*Mutator* foci intensity varies along the adult germline. (A) Fixed *mut-16::gfp::3xFLAG* adult germlines were immunostained for MUT-16 (anti-FLAG, green), SYP-1 (anti-SYP-1, red) and DNA (blue) to visualize *Mutator* foci intensity pattern along the germline in relation to mitotic or meiotic cell state. (B) Graph plotting percentage of above-threshold foci along 10 equal sections of *mut-16::gfp::3xFLAG* germlines. n=3. (C) *mut-16::gfp::3xFLAG; him-8(tm611)* mutants display an extended transition zone (white dashed line) indicated by DAPI-visualized transition nuclei throughout gonad (white arrows). MUT-16 foci (anti-GFP, green) intensity pattern remains similar to the wild-type. (D) *chk-2* RNAi disrupts polarization of DAPI-visualized transition zone nuclei (yellow dashed line). MUT-16 foci (anti-FLAG, green) intensity pattern remains similar to the wild-type. (E) *gld-1 gld-2* RNAi produces a tumorous mitotic germline. MUT-16 foci (anti-FLAG, green) intensity is bright within the proximal mitotic tumor (asterisks). (F) Quantification of percent above-threshold foci in *gld-1 gld-2* RNAi gonads. n= 4. (G) *atx-2* RNAi germline MUT-16 foci (anti-FLAG, green) intensity is more evenly distributed than wild-type foci intensity. (H) Quantification of percent above-threshold foci in *atx-2* RNAi gonads. n= 4. Distal gonad tips are oriented to the left in all images. In all indicated images, SYP-1 (red) is used as a marker for meiotic progression and DNA is visualized with DAPI (blue). All scale bars, 15μm.

### *Mutator* foci intensity is associated with mitotic and not transition zone nuclei

Because the number of above-threshold *Mutator* foci peaks near the transition zone, we sought to determine if foci intensity was associated with the polarization of chromosomes that produces the unique crescent-shaped morphology of transition zone nuclei. To first examine the effects of an extended transition zone, we utilized a mutation in the *him-8* gene, required to promote X chromosome pairing and synapsis (Phillips *et al*. 2005). Cells that fail to complete synapsis cannot exit the condensed transition state, resulting in polarized nuclei well into the pachytene region of the germline. Despite having an extended transition zone region, the brightest *Mutator* foci in a *him-8* mutant gonad remained restricted to the mitotic region and beginning of the transition zone, with an overall pattern similar to wild-type animals (Figure 2C). Next, we aimed to determine if *Mutator* foci intensity was affected by eliminating the polarized nuclear morphology in the transition zone. CHK-2, an ortholog of human checkpoint kinase 2, is required both for homolog pairing and spatial reorganization of chromosomes at the onset of pairing; *chk-2* mutant germlines do not contain any polarized transition zone nuclei (MacQueen and Villeneuve 2001). We knocked down *chk-2* by RNAi and found that, though *chk-2* RNAi gonads lacked polarized nuclei, the intensity pattern of *Mutator* foci remained similar to wild-type, with a peak in intensity at the mitotic to meiotic transition (Figure 5D). Our data indicates that neither extension nor morphological disruption of the transition zone nuclei perturbs the intensity pattern of *Mutator* foci.

Because the mitotic region is associated with some of the highest numbers of above-threshold *Mutator* foci, we next sought to investigate whether the mitotic cell state determines *Mutator* foci intensity. To this end, we knocked down the KH motif-containing RNA-binding protein GLD-1 and the poly(A) polymerase GLD-2, which are redundantly required to promote meiotic entry and, when disrupted, produce a tumorous mitotic germline (Kadyk and Kimble, 1998). Because the RNAi knockdown is not completely penetrant, we observe some nuclei entering meiosis, marked by SYP-1 staining (Figure 5E). Despite incomplete penetrance, the proximal region of the germline produces a large mitotic tumor which contains very bright *Mutator* foci. Quantification reveals that significantly more above-threshold foci are found in the proximal tumor than even the distal mitotic zone. Additionally, no peak of fluorescence is found at the onset of SYP-1 loading, in contrast with the peak near wild-type transition zones (Figure 5F). In gonads where the RNAi is more penetrant and produces a larger mitotic tumor, bright *Mutator* foci are found more uniformly throughout the germline (Supplemental Figure 4). These observations indicate that the mitotic cell state is a contributing factor of *Mutator* foci intensity.

Finally, we sought to manipulate germline morphology and cell state in additional ways to test if *Mutator* foci intensity experienced similar perturbations. We knocked down ATX-2, which functions to promote germ cell proliferation and prevent premature meiotic entry; RNAi of *atx-2* was previously observed to produce small germlines with truncated mitotic and transition zones (Maine *et al*. 2004). As expected, immunostained gonads were smaller than wild-type and had reduced mitotic tip and transition zones (Figure 5G). Interestingly, *Mutator* foci appeared bright in the mitotic tip but did not form a distinct intensity peak preceding the transition zone and, in the ensuing pachytene region, foci generally appeared brighter than in the mid-pachytene of wild-type animals. Quantification reveals that above-threshold foci are more evenly distributed throughout the germline (Figure 5H). Thus, distribution of above-threshold *Mutator* foci can also be altered by general perturbations of germline morphology and cell state.

## DISCUSSION

*Mutator* foci are hubs of secondary siRNA biogenesis in the *C. elegans* germline. Here we show that *Mutator* foci first appear in PGCs beginning around the 100-cell stage of embryogenesis, and that these foci persist in germ cells through all subsequent developmental stages. Additionally, we find MUT-16 present in post-meiotic spermatids. We further demonstrate that both the presence of RNA and the germline cell cycle play key roles in promoting assembly of *Mutator* foci. Therefore, our work begins to define both the developmental stages and regulatory factors that shape *Mutator* foci integrity and intensity.

### *Mutator* foci in embryos

Our first observation of robust *Mutator* foci occurs around the 100-cell stage, after the P4 cell divides into the PGCs Z2 and Z3. Interestingly, this timing coincides with the appearance of Z granules, which are germline-specific phase-separated condensates required for RNAi inheritance. At this time, Z granules de-mix from P granules to form adjacent foci (Wan *et al*. 2018). MUT-16, however, does not appear significantly enriched in P granules prior to *Mutator* foci formation and is therefore unlikely to be de-mixing from P granules, but rather forming foci *de novo*. The birth of the Z2 and Z3 PGCs is also marked by the disappearance of MEG-3 and MEG4, which surrounds the PGL phase of P granules in early embryogenesis and are crucial for P granule assembly (Wang *et al*. 2014). Together, these data suggest that there may be a coordinated reorganization of germ granule components at this time. While the exact mechanism for this reorganization remains unknown, the timing coincides with a burst of transcription in the PGCs (Seydoux and Dunn 1997). It is possible that an increase in *mut-16* transcript, and thus protein levels, leads to the emergence of *Mutator* foci; however, detectable levels of cytoplasmic MUT-16 protein can be observed in all cells throughout embryonic development. Therefore, we favor the possibility that newly synthesized transcripts emerging from the nuclear pores necessitate and perhaps directly promote assembly of the P granule/Z granule/*Mutator* foci multi-condensate structures.

### Regulation of *Mutator* foci by RNA

Because the appearance of *Mutator* foci in embryos coincides with the onset of transcription in the PGCs, we investigated the effect of transcriptional inhibition in the adult germline. It was previously shown that injection of a transcriptional inhibitor causes pachytene-specific loss of P granules (Sheth *et al*. 2010). We found that *Mutator* foci similarly rely on RNA for their stability in the mid-pachytene region. However, in late pachytene we observed differences in the dissolution of *Mutator* foci and P granules following transcriptional inhibition. Specifically, we observed persistent *Mutator* foci in the late pachytene, despite visible dissipation of all P granules at seven hours post-transcriptional inhibition. The sustained presence of *Mutator* foci in late pachytene, but not mid pachytene, could arise for multiple reasons. One possibility is that *Mutator* foci may be interacting with more stable classes of RNAs. A recent discovery shows that long stretches of alternating uridine (U) and guanosine (G), termed polyUG (pUG) tails, are added to siRNA-targeted transcripts to mark them as templates for secondary siRNA synthesis (Shukla *et al*. 2020). These pUG tails, added by the *Mutator* complex protein MUT-2, are hypothesized to confer stability to targeted transcripts. If these stabilized transcripts are associating with *Mutator* foci in higher quantities in late pachytene, it may explain the resistance of late pachytene *Mutator* foci to dissolution after transcriptional inhibition. An alternate explanation may reside in the phase-separation properties of *Mutator* foci. We previously tested Fluorescence Recovery After Photobleaching on *Mutator* foci in the late-pachytene region, and showed that, although there was rapid recovery of a highly-mobile fraction within *Mutator* foci, there was also a large non-mobile fraction of MUT-16 that failed to recover after photobleaching. We have not yet identified the molecular basis of these distinct phases within *Mutator* foci, but phase separated granules can be regulated by post-translational modifications (Li *et al*. 2013; Wang *et al*. 2014), making this a tantalizing avenue for future exploration. Nonetheless, this non-mobile fraction may be in a more gel-like state and therefore resistant to dissipation after transcriptional inhibition. A direct comparison of the liquid properties of mid versus late-pachytene *Mutator* foci is necessary to address this possibility.

We also noticed that P granules became larger and more rounded at three hours post-transcriptional inhibition in the late pachytene compared to mid-pachytene. Typically, P granules are intimately associated with nuclear pores, likely through the interaction of the FG-repeats in DEAD-box helicases GLH-1, GLH-2, and GLH-4, with nuclear pore-containing FG-repeat proteins (Updike *et al*. 2011; Marnik *et al*. 2019). This non-spherical appearance of P granules due to association with nuclear pores has been described as a “wetting” of the nuclear membrane (Brangwynne *et al*. 2009). Interestingly, a mutation in GLH-1 also creates large, round P granules, possibly caused by an inability to release captured RNA (Marnik *et al*. 2019). Thus, our data suggests that P granule wetting may also depend on the sustained presence of transcripts exiting the nucleus.

### *Mutator* foci in the cell cycle and inheritance

Finally, we explored why *Mutator* foci brightness varies within different regions of the germline and determined that perturbations to the cell cycle affected the intensity and distribution of *Mutator* foci. We found that nuclei in mitosis tend to be associated with large, bright *Mutator* foci, whereas *Mutator* foci in the early and mid-pachytene are much dimmer. The exact mechanism governing *Mutator* foci robustness remains elusive. Interestingly, the mid-pachytene region is also where *Mutator* foci are most sensitive to transcriptional inhibition, again suggesting that more stable RNAs, or perhaps higher concentrations of small RNA-target transcripts, in the mitotic and late pachytene regions may promote condensation of larger foci.

It has been well-documented that small RNAs can be inherited through both the maternal and paternal germline (Grishok *et al*. 2000; Alcazar *et al*. 2008; Lev *et al*. 2019). This transgenerational epigenetic inheritance can promote a memory of self and non-self transcripts across generations as well as transmit gene regulatory information in response to environmental conditions (Ashe *et al*. 2012; Luteijn *et al*. 2012; Shirayama *et al*. 2012; Conine *et al*. 2013; Rechavi *et al*. 2014; de Albuquerque *et al*. 2015; Phillips *et al*. 2015). While the Argonaute WAGO-4 and helicase ZNFX-1 have been implicated in the transmission of maternal small RNAs (Ishidate *et al*. 2018; Wan *et al*. 2018; Xu *et al*. 2018), little is known about how small RNAs are packaged and transmitted through paternal lineages. Here, we observed *Mutator* foci throughout spermatogenesis and detected MUT-16 in the cytoplasm of post-meiotic spermatids. A similarly granular expression pattern in spermatids was seen for WAGO-1 (Conine *et al*. 2010); if these granules coincide, further work will be necessary to determine if they act together to promote paternal inheritance and whether they promote paternal deposition of not just small RNAs, but also small-RNA targeted mRNAs. Ultimately, *Mutator* foci morphology and regulation may influence efficacy of RNAi in certain cell stages or environments, an avenue that warrants further investigation.

## ACKNOWLEDGEMENTS

We thank the members of the Phillips lab for helpful discussions and feedback on the manuscript, and the Updike lab for granularity quantification protocols. This work was supported by the National Institute of Health grants R35 GM119656 (to C.M.P.) and T32 GM118289 (to D.C.W.), and the National Science Foundation Graduate Research Fellowship Program Grant No. DGE 1418060 (to C.J.U.). CMP is a Pew Scholar in the Biomedical Sciences supported by the Pew Charitable Trusts (www.pewtrusts.org) and C.J.U. is a USC Dornsife-funded Chemistry-Biology Interface trainee. D.A. is a student at Downtown Magnets High School and performed research under the mentorship of C.J.U. The *him-8(tm611)* allele was isolated and provided by Shohei Mitani and the Japanese National BioResource for *C. elegans*. The funders had no role in study design, data collection and analysis, decision to publish, or preparation of the manuscript.

---

**Supplemental Figure 1.**
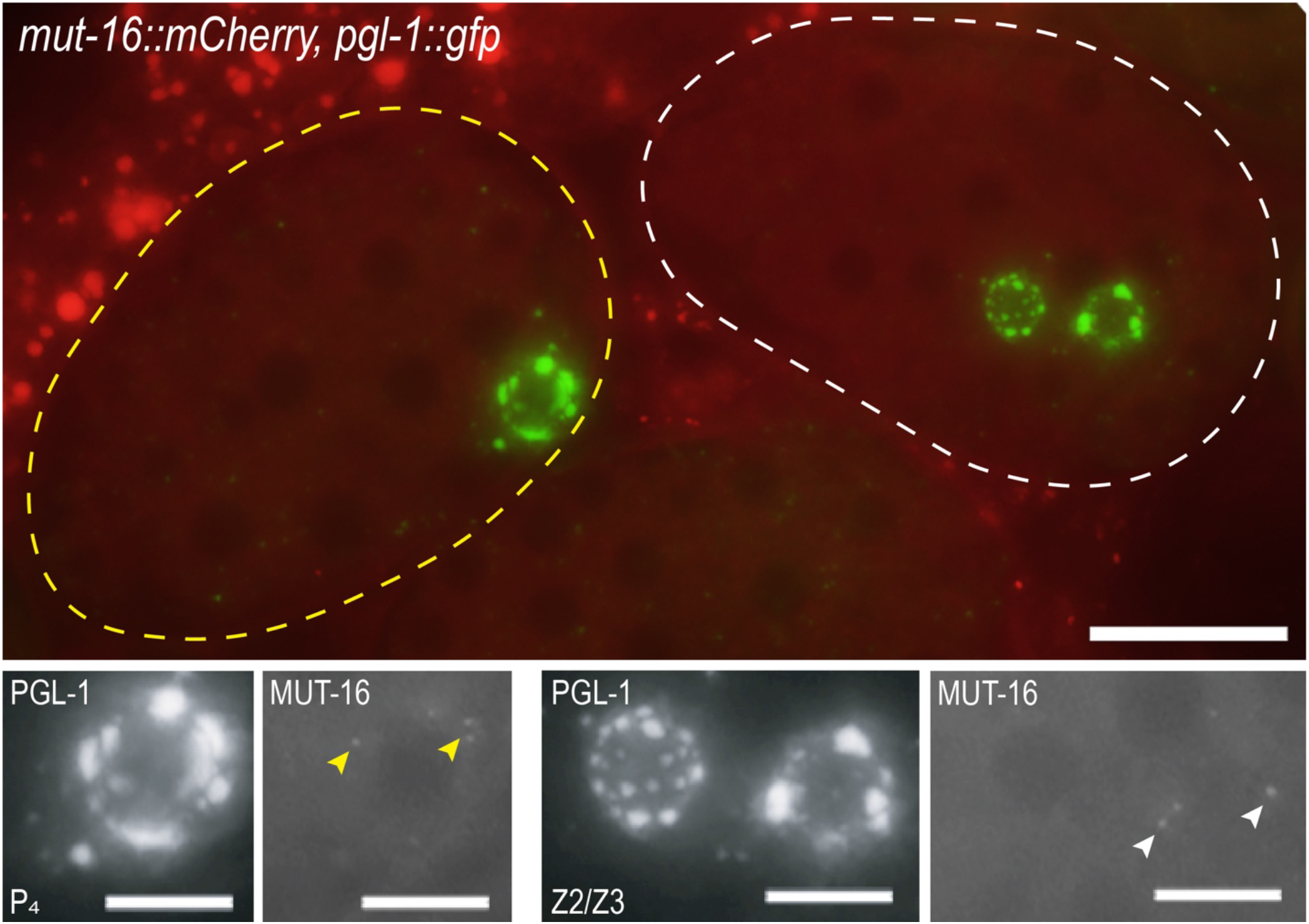
*Mutator* foci are weakly present in the 30-cell stage. Live image of a 30-cell embryo (yellow outline) and 100-cell embryo (white outline) within an undissected *mut-16::mCherry; pgl-1::gfp* adult. PGL-1::GFP (green) granules associate with the P4 and Z2/Z3 progenitor germ cells (insets). Small MUT-16::mCherry (red) puncta are visible in the P4 cell of the 30-cell embryo (yellow arrowheads). MUT-16 foci are clearly visible in the 100-cell stage (white arrowheads). Image is a maximum intensity projection from five Z-stacks. Top scale bar, 15μm. Bottom scale bars, 5μm.

**Supplemental Figure 2.**
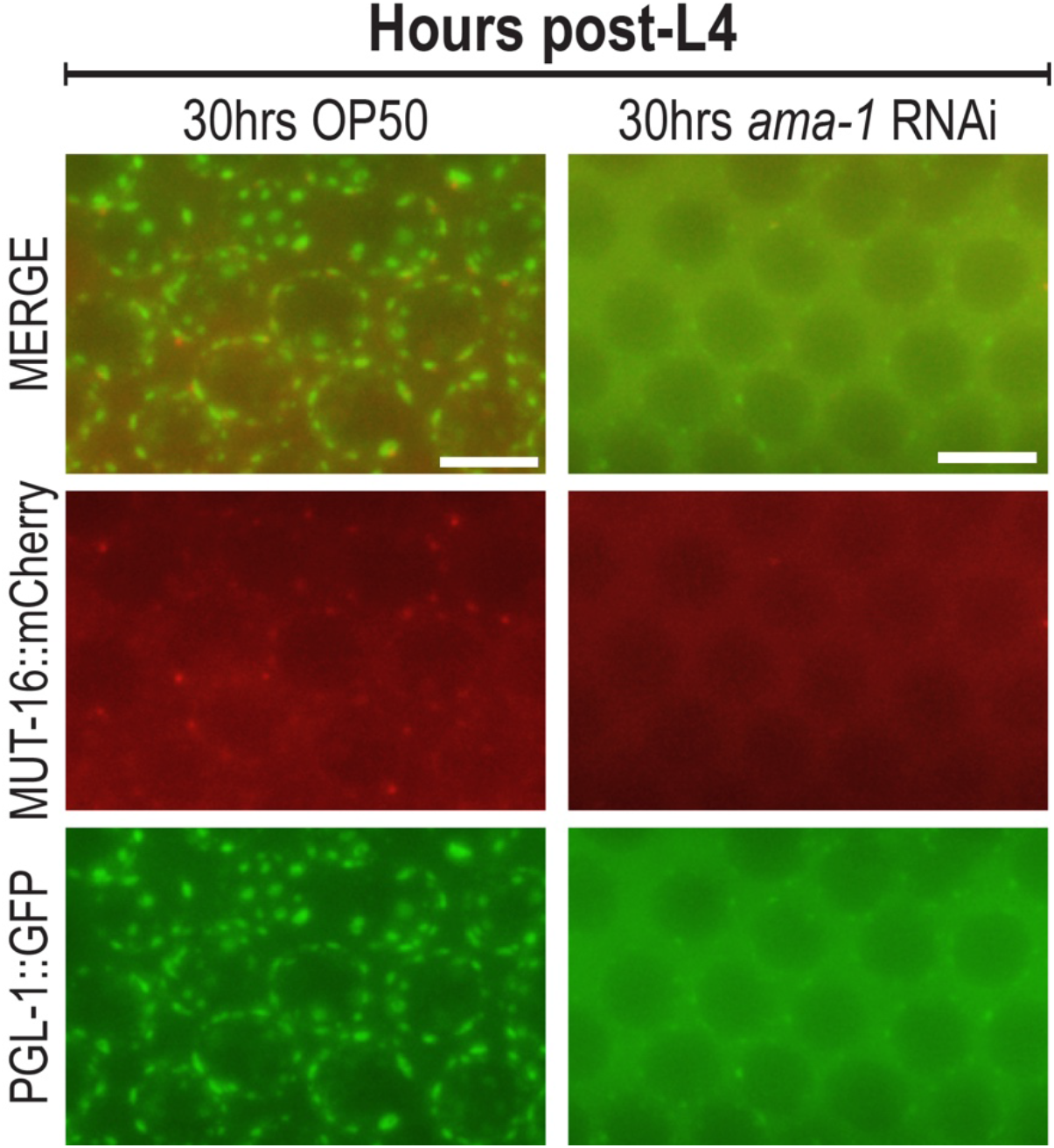
*Mutator* foci dissipate after *ama-1* RNAi. (A) Larval L4 *mut-16::mCherry; pgl-1::gfp* animals were placed on OP50 or on *ama-1* RNAi for 30 hours. MUT-16 foci and PGL-1 granules are present in the mid-pachytene after 30 hours post-L4 on OP50, but dissipate after 30 hours post-L4 on *ama-1* RNAi. Images are maximum intensity projection of five Z-stacks. Scale bars, 5μm.

**Supplemental Figure 3.**
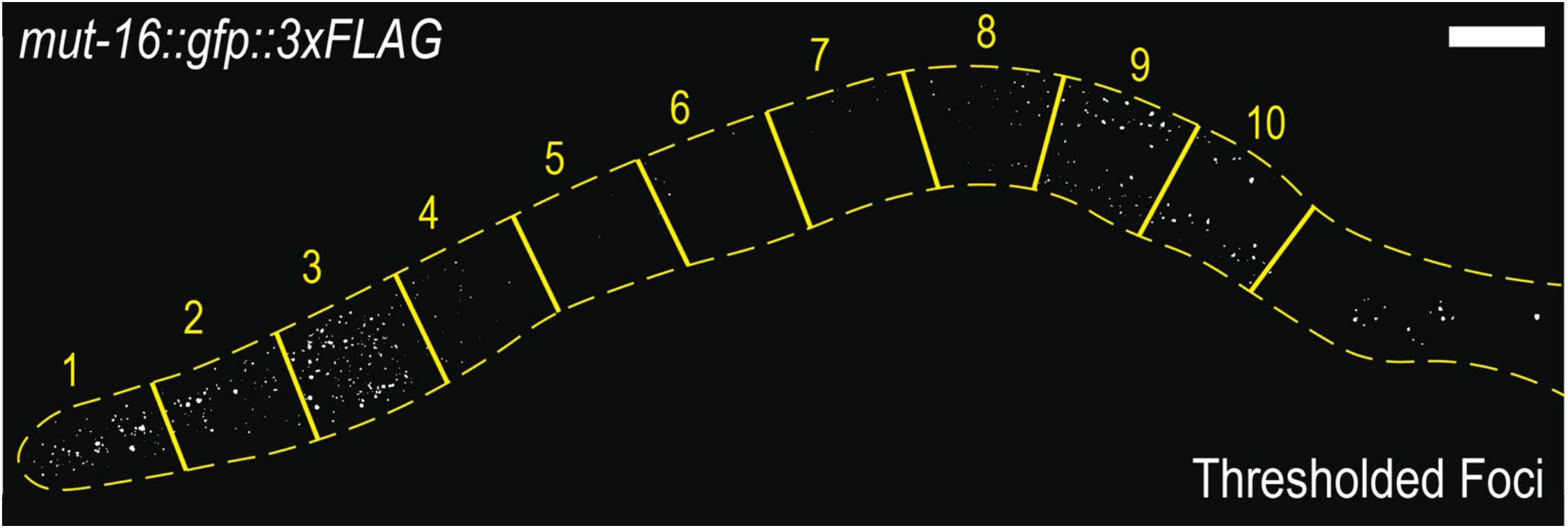
Image thresholding reveals bright *Mutator* foci. Gonads were divided by length into 10 equal regions ending at the first single-file diplotene/diakinesis nucleus (yellow borders). After identical thresholding, images displayed only above-threshold foci (white dots) which were quantified using the analyze particles function in FIJI.

**Supplemental Figure 4.**
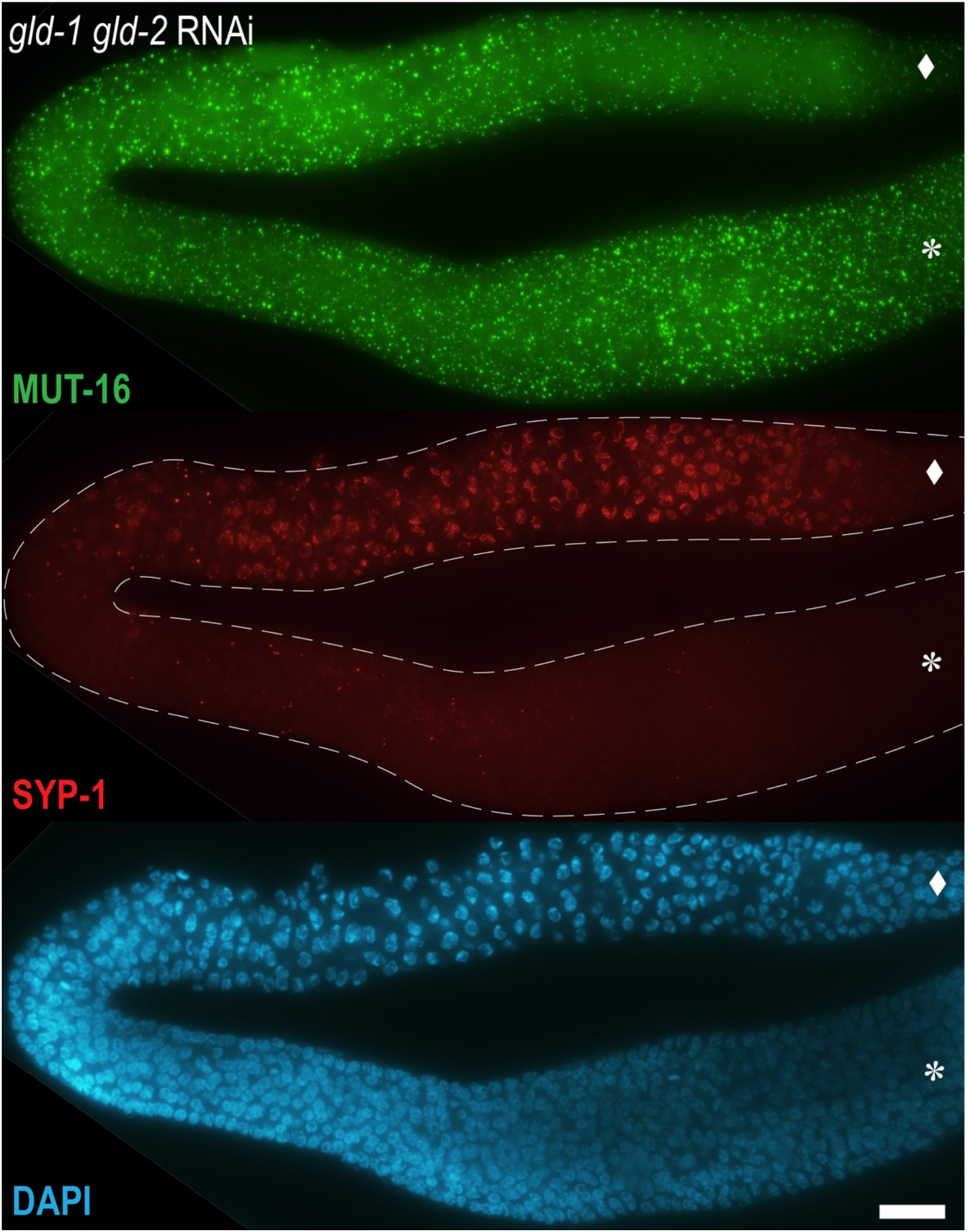
*Mutator foci* are bright in the mitotic tumor of *gld-1 gld-2* RNAi gonads. Double *gld-1 gld-2* RNAi on *mut-16::gfp::3xFLAG* animals produces a large proximal mitotic tumor (asterisks). Top: Numerous bright MUT-16 foci (anti-FLAG, green) are found throughout the mitotic tumor. Middle: Due to incomplete penetration, some SYP-1 (anti-SYP-1, red) staining is present in the more distal region (diamond). The proximal tumor lacks SYP-1 staining, indicating cells in mitosis. Bottom: DAPI-visualized DNA (blue) reveals an abundance of small mitotic nuclei in the proximal tumor. Note that the complete distal tip of the gonad is not shown. Images are a maximum intensity projection of 25 Z-stacks. Scale bar, 15μm.

## Notes

### Competing Interest Statement

The authors have declared no competing interest.

